# Hybotidae (Diptera) of the Botanic Garden Jean Massart (Brussels-Capital Region, Belgium) with description of two new *Platypalpus* species and comments on the Red Data List

**DOI:** 10.1101/2022.12.26.521931

**Authors:** Patrick Grootaert

## Abstract

Ninety hybotid species are recognized in the Botanic Garden Jean Massart (Brussels), representing 52 % of the hybotids ever recorded in Belgium. Two species new to science are described: *Platypalpus massarti* sp. nov. and *P. pictitarsoides* sp. nov. Following species are reported for the first time in Belgium: *Drapetis infitialis* (Collin, 1961), *Platypalpus negrobovi* Grootaert, Kustov & Shamshev, 2012 and *Trichina opaca* Loew, 1864. In addition, comments are given on a selected number of species: *Bicellaria intermedia* Lundbeck, 1910, *Platypalpus aurantiacus* (Collin, 1926), *Platypalpus longimanus* (Corti, 1907), *Platypalpus nanus* (Oldenberg, 1924), *Platypalpus rapidoides* Chvála, 1975, *Platypalpus subtilis* (Collin, 1926), *Stilpon subnubilus* Chvála, 1988 and on the genus *Hybos* Meigen, 1803. The holotype of *P. cryptospina* (Frey, 1909) is revised.

Only 30 species or 33% of the species present are in a ‘Safe/Low risk’ Red Data Book category meaning that the other 66% are in a more or less ‘Threatened’ category.

## Introduction

The present study reports on Hybotidae examined as part of a comprehensive three-year survey of the Diptera in the Botanic Garden Jean Massart (Oudergem, Brussels-Capital Region, Belgium). This tiny botanic garden of 4.5 ha, is squeezed in between the eastern border of the city of Brussels and the Sonian forest. Nearly 2,000 plant species have been recorded in this Natura 2000 site. The area is composed of various biotopes such as humid areas with a swamp and ponds, an old orchard on dry grassland, a medicinal plants garden, an arboretum and an evolution garden. All is mixed with patches of semi natural woods.

In a previous study on the genus *Drapetis* Macquart, five species were found, including *Drapetis bruscellensis* Grootaert, 2016, a new species for science. In the present paper attention is paid to all the hybotid genera and species of special interest are commented. The Red Data Book status of the various hybotid species was examined (Grootaert *et al*., 2001) in order to assess the value of the biodiversity in the Botanic Garden Jean Massart.

## Material and methods

The present survey is based on a sampling in six sites with Malaise traps. However, the sampling in the various sites was spread over three years as is shown in Table 1. The present study is limited to Malaise trap sampling only.

**Table 1.** Overview of the sampling effort per site and a summary of the habitat characteristics.

|  | MT1 | MT2 | MT3 | MT4 | MT5 | MT6 |
| --- | --- | --- | --- | --- | --- | --- |
|  | 50°48'52.57"N<br>4°26'20.31"E | 50°48'52.70"N<br>4°26'22.50"E | 50°48'50.68"N<br>4°26'22.14"E | 50°48'52.83"N<br>4°26'15.62"E | 50°48'47.12"N<br>4°26'15.90"E | 50°48'49.02"N<br>4°26'6.97"E |
| start | 7.V.2015 | 7.V.2015 | 21.IV.2016 | 14-III-2017 | 14-III-2017 | 14-III-2017 |
| end | 23.III.2017 | 23.III.2018 | 14.III.2017 | 23.III.2018 | 23.III.2018 | 23.III.2018 |
| years | 2 | 3 | 1 | 1 | 1 | 1 |
| soil | humid | humid | humid | dry | dry | swampy |
|  | loamy | loamy | loamy | loamy | loamy | loamy-sand |
| exposure | partly shaded | sun exposed | shaded | shaded | sun exposed | partly shaded |
| biotope | border of meadow,<br>behind compost heap | border of flowerbeds,<br>border of hedge | inside small wood; next to heap of decomposing wood | arboretum; no or short herbaceous layer | flowered meadow in old apple orchard | close to reed bed, above small ponds |

The hybotid material was collected and stored in ethanol. In the listing of the records ‘reg.’ refers to the register number of the specimens of the Diptera from the Botanic Garden Jean Massart in the collections of the Royal Belgian Institute of Natural Sciences in Brussels (RBINS).

### Analysis Red Data Book Data

The Red Data Book (RDB) categories used in the present study are those listed in Grootaert *et al*. (2001) for the Empididae *sensu lato*. At that time the empidoids (minus the Dolichopodidae) contained the Empididae, Hybotidae, Microphoridae (now as Microphorinae in the Dolichopodidae), Atelestidae and some Brachystomatidae. In this study, all species were assigned to Red Data Book categories which are based on a combination of a rarity and a trend criterion. Rarity is expressed as the proportion of the total number of UTM 5 km squares sampled in which the species have been found since 1981. The trend criterion is interpreted as the change of the species rarity between 1887-1980 and 1981-1999. A comparable number of UTM 5 km squares was investigated during the two time periods. In addition, threatening of the specific habitat where the species is living was also taken into account as a criterion for the categories. Since records from Brussels-Capital Region in which the Botanic Garden Jean Massart is situated, were included in the Red Data Book of Flanders, the RDB can be used for an assessment of the empidoids in the Botanic Garden Jean Massart.

## Observations

All the hybotid flies (except those of a few missing samples) were identified that were collected during the three-year survey (from May 2015 until April 2018) at the Botanic Garden Jean Massart. This resulted in 4,192 specimens belonging to 90 species.

Annexe 1 shows an overview of the species and their occurrence in the six investigated sites (MT1 to MT6). Since site 1 (MT1) and site 2 (MT2) were respectively sampled during 2 and 3 years, the data are presented for each year separately (MT1-2015, MT1-2016; MT1-2015, MT1-2016 and MT1-2017) which allows a quick overview of the differences per year. This separation per year also illustrates the turnover of the species for each season. This is best illustrated in site 2 (MT2) where a total number of 79 species were recorded over the three-year survey. The turnover of the number of species is large: 58 species were recorded in 2015, 61 species in 2016 and 59 species in 2017. Although the number of species per year is more or less the same in MT2, the turnover per year is very high: in 2016 there were 12 species absent from 2015, but 15 were new; in 2017 there were 18 species not present in 2015 and 17 species not present in 2016.

In addition to the differences in turnover per year, there is a difference of diversity in site 1 (MT1) and site 2 (MT2), laying just 43 m opposite to each other in the evolution garden. In site 1, 41 species were recorded in 2015 while 42 species in 2016. In site 2, 58 species were recorded in 2015 and 61 species in 2016. This difference in diversity is attributed to the different insolation of the two sites. Site 1 receives direct sunlight in early morning only, while site 2 receives direct sunlight from noon until late evening. Likely, site 2 is warming up more than site 1 so that the activity of the flies is different which is reflected not only in the number of specimens sampled but also in the number of species.

Site 3 (MT3) is continuously in the shade, with an undergrowth of grasses and a huge pile of decaying wood. Only 38 hybotid species were found in this site but the largest population of *Platypalpus optivus* in the Garden was observed in this site. The population of *Platypalpus exilis* and *P. luteolus*, both yellow species, was also the largest at this site of the Garden. This is not exceptional, since these yellow species thrive in shaded conditions. On the other hand, the population of the yellow *Elaphropeza ephippiata* was very low although grasses were ample present in the undergrowth being a favourite microhabitat of *E. ephippiata*.

Site 4 (MT4) has the lowest diversity of all sites. Only 57 specimens were found belonging to 24 species. The arboretum is characterised by a partly naked soil or a very low herb layer from spring onwards. There is never direct sunlight.

Site 5 (MT5) has 44 species. In this old apple orchard, *P. aristatus* was dominant in spring and several other ubiquist species were abundant such as *P. longicornis, P. pallidiventris* and *P. calceatus* amongst others.

In Site 6 (MT6) *Platypalpus pictitarsoides* sp. nov. was the dominant species with 123 specimens. It was also the site were most specimens of this species were found.

## Annotated checklist with description of new species

### *Bicellaria intermedia* Lundbeck, 1910

#### Material examined

Oudergem, Botanic Garden Jean Massart: **MT2**, 1♂, 1-8.VII.2015.

Additional material examined: Belgium: 1♀, Parette (prov. Luxembourg), (31UFR91), 3.VII.1980; 1♂, Vecquée (32UKB90), 10.VII.1951 (R. Tollet); 1♂, 1♀, Bihain (prov. Luxembourg), (31UGR06), 6.VI.1952 (R. Tollet); 2♂♂, Fagne des Mochettes, Samrée (31UFR96), 5.VI.1952 (R. Tollet); 5♂♂, 2♀♀, Samrée (31UFR86), 5.VI.1952 (R. Tollet); 6♂♂, 3 ♀♀, Franc Bois (31UFR33), 18.VI.1958 (leg. A Collart).

*Bicellaria intermedia* was not yet recorded from Flanders nor Brussels Region and is hence not in the Red Data Book (Grootaert *et al*., 2001), however several specimens were recorded earlier from southern Belgium.

### *Drapetis infitialis* (Collin, 1961)

#### Material examined

Oudergem, Botanic Garden Jean Massart: **MT2**: 2♂♂, 8-15.VI. 2017 (reg. 1364); **MT5**: 1♂, 22-30.VI.2017 (reg. 1130); 1♂, 19-26.VII.2017 (reg.1161).

This species which is closely related to *D. exilis*, is new for the Belgian fauna. It differs from the latter species mainly by the large tip of the right cercus (Collin, 1961).

## Genus *Hybos* Meigen, 1803

### *Hybos culiciformis* (Fabricius, 1775)

#### Material examined

Oudergem, Botanic Garden Jean Massart: **MT1**: 3♀♀, 13-27.X.2016 (ref. 571); **MT2**: 1♀, 26.VI-1.VII.2015 (ref. 158); 1♀, 14-28.VII.2016 (ref. 920); 1♂, 23.VIII-1.IX.2017 (ref. 1352).

### *Hybos femoratus* (Müller, 1776)

#### Material examined

Oudergem, Botanic Garden Jean Massart: **MT2**: 1♀, 22.IX-12.X.2017.

During the three consecutive years of the survey, only six specimens of *Hybos culiciformis* and one specimen of *Hybos femoratus* were recorded.

*Hybos culiciformis* is considered as fairly common but ‘Near threatened’ due to a significant decline since 1981 (Grootaert *et al*., 2001). *Hybos femoratus* is considered as ‘Vulnerable’ and considered as fairly ‘Rare’ with a strong decline of the populations. The third Belgian species *H. grossipes*, quoted as ‘Endangered’ with a significant decline of the populations, was not found at all during the present survey.

*Hybos* species are predators that catch small insects in the air with their huge raptorial hind legs. Although there is a very high diversity of microhabitats with scrubs and vegetation that can be used as look-out for *Hybos* and moreover ample prey is available, the abundance is very low. It seems that the general trend of decline that was already mentioned in Grootaert *et al*. (2001) is continuing.

## Genus *Platypalpus*

*Platypalpus* species are small predators that catch prey landing on leaves of the vegetation. Forty-nine species were recorded here in the Botanic Garden Jean Massart.

### *Platypalpus aurantiacus* (Collin, 1926)

*Tachydromia aurantiaca* Collin, 1926: 152.

*Tachydromia aurantiaca* in Collin, 1961: 207, description.

*Platypalpus aurantiacus* (Collin, 1926) in SMITH & CHVáLA, 1976: 139, illustration male terminalia.

*Platypalpus aurantiacus* (Collin, 1926) in Chvála, 1989: 279, diagnosis and illustration male antenna (Fig. 12).

#### Material examined

Oudergem, Botanic Garden Jean Massart: **MT1**: 1♀, 3-9.VI.2016 (ref. 418); 1♀, 6-11.V.2016 (ref. 596); **MT2**: 1♀, 28.V-4.VI.2015 (ref. 578); 1♂, 21-28.V.2015 (ref. 584); 1♀, 21-28.V.2015 (ref. 627); 2♀♀, 13-21.V.2015 (ref. 686); 1♀, 13-21-May 2015 (ref. 745); 1♂, 6-11.V.2016 (ref. 1311); 1♀, 5-11.V.2017; 2♀♀, 17-24.V.2017 (ref. 1256); 1♂, 1-8.VI.2017 (ref. 1268); **MT3**: 1♀, 3-9.VI.2016 (ref. 445); **MT4**: 1♂, 24.V-1.VI.2017 (ref. 1087); **MT5**: 1♀, 11-17.V.2017.

Additional material examined: Belgium: 2♀♀, Buzenol, 19.V.1981; 5♀♀, Buzenol, 2.VI.1981 (leg. P. Grootaert).

#### Comments

This rare species was previously only known from the extreme South of Belgium where it was collected in a Malaise trap at the border of a deciduous forest. Here in the Botanic Garden it was found in five of the six investigated sites, always in very low numbers except in site 2 (MT2) where over the three years of sampling 11 specimens were recorded. It was not found in the marshland (MT6).

#### Distribution

According to the Fauna Europaea it is recorded in Austria, Belgium, British Isles, Czech Republic, France, Germany and Hungary.

### *Platypalpus massarti* sp. nov

Figs 1–2

**Fig. 1.**
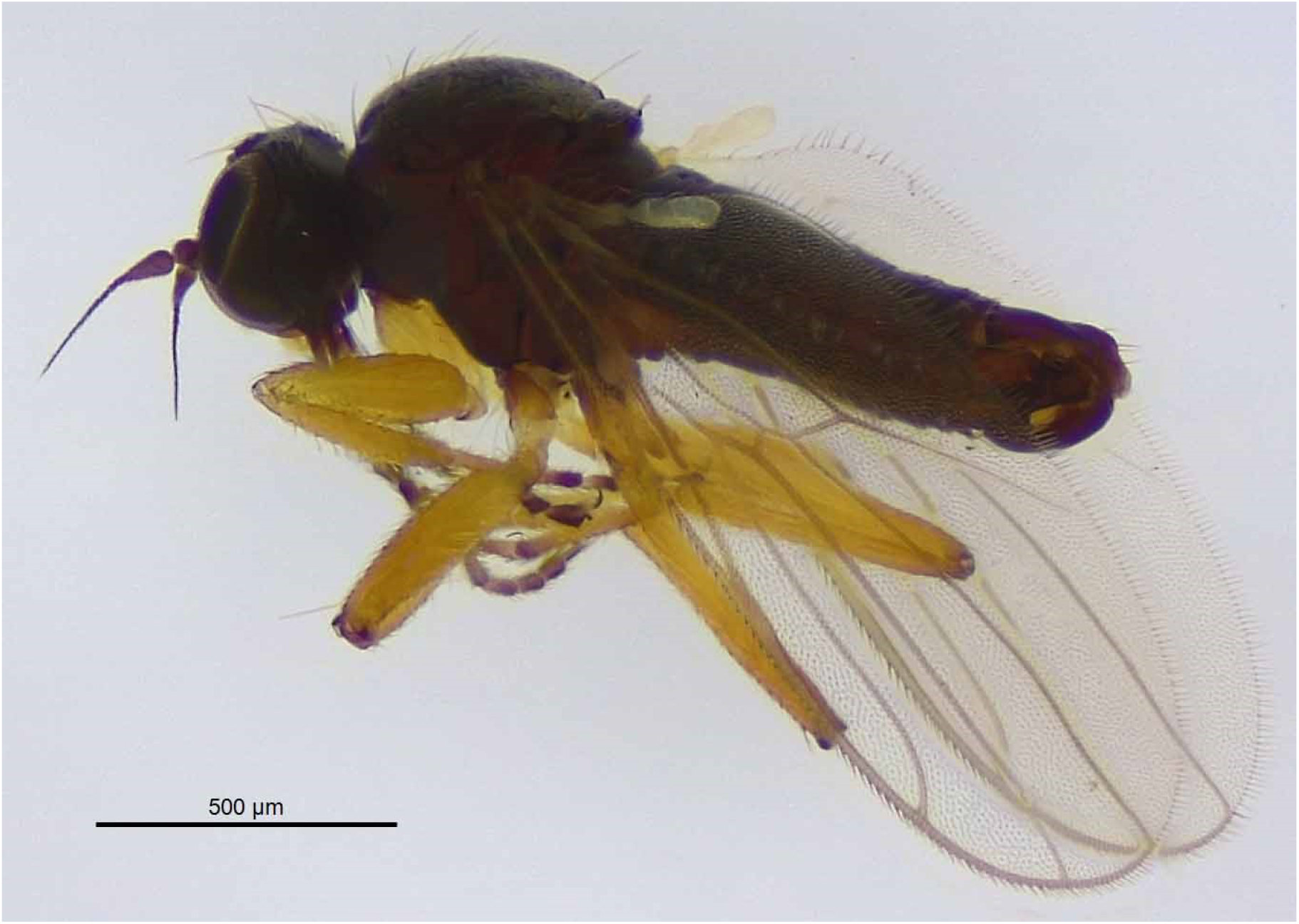
*Platypalpus massarti* sp. nov., holotype male, habitus (photo Isabella Van de Velde).

**Fig. 2.**
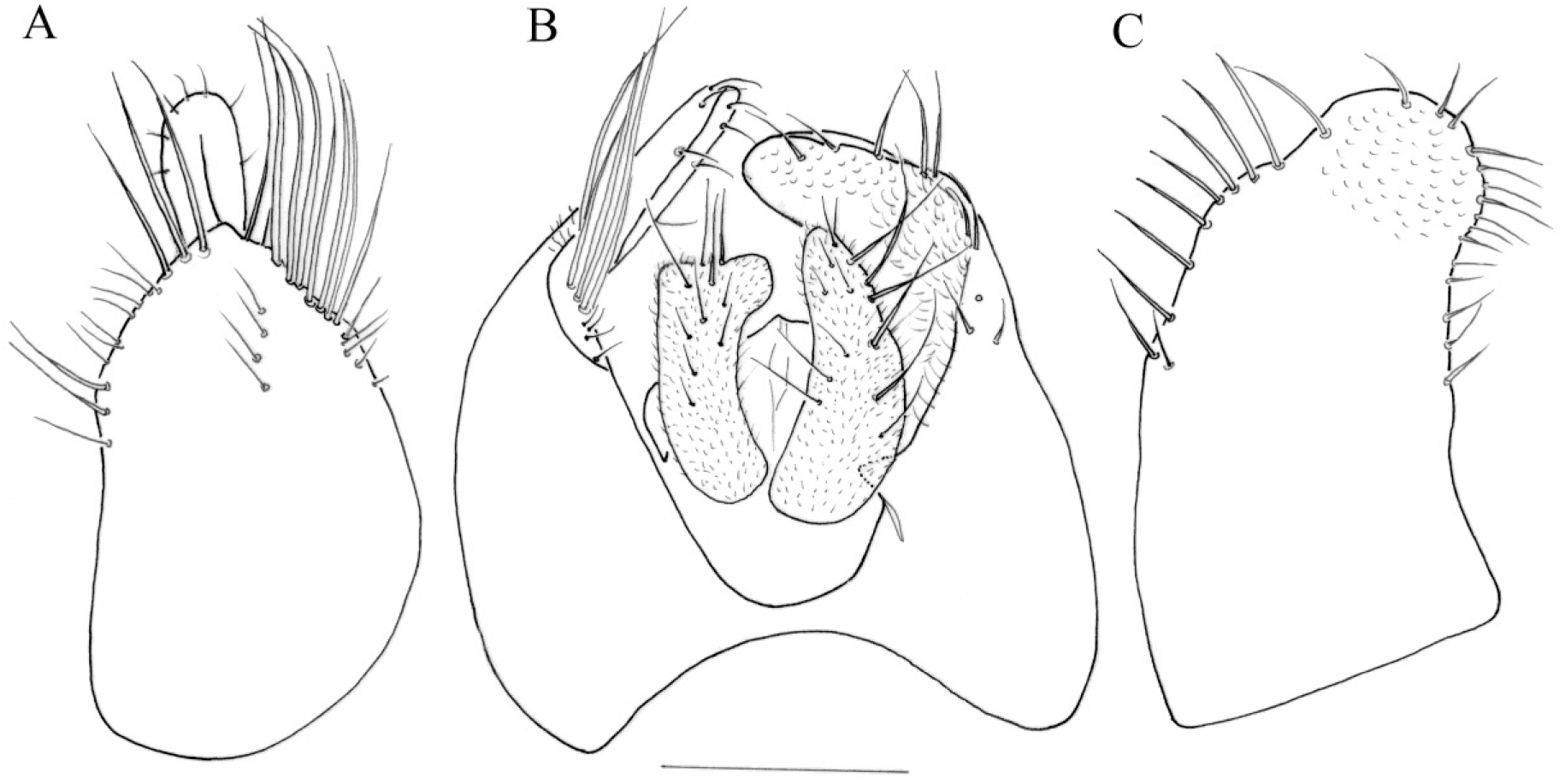
*Platypalpus massarti* sp. nov., holotype male, terminalia. A, right epandrial lamella. B, epandrium dorsal. C, left epandrial lamella. Scale 0.1 mm.

urn:lsid:zoobank.org:act:942C978F-D0E4-4B0D-B633-D5C34F6A7A6A

#### Material examined

Holotype ♂: Belgium, Oudergem, (Brussels) Botanic Garden Jean Massart, **MT2**, 8-15.VI.2017 (ref. 1377, RBINS); Paratype: Belgium, Oudergem, (Brussels) Botanic Garden Jean Massart, 1 ♀, **MT2**, 23.VI-1.VII.2016 (ref. 872, RBINS).

#### Diagnosis

A small black species (1.5-1.6 mm) with one pair of yellow vertical bristles. Antenna entirely black. Postpedicel in male 2.5 times as long as wide, in female 2 times; arista 2 times as long as postpedicel. Palpus pale yellow with long pale apical bristle. Mesoscutum dusted. Sternopleura with a shiny black patch. Acrostichals short, biserial and widely separated. Four long dorsocentrals. Legs yellow, except for mid and hind coxae brown and all legs with all tarsomeres brown annulated, apical two tarsomere entirely brown. In female apical annulation of basal tarsomeres shorter and less pronounced than in male. Fore tibia in male only weakly swollen. Mid femur with a few short pale posteroventral bristles. Mid tibia in male and female with a very short, triangular, pale brownish apical spur. Left epandrial lamella with only short marginal bristles on the left side, limited to the apical half.

#### Etymology

The new species is dedicated to Prof Jean Massart, ecologist avant-la-lettre, who created 100 years ago the Botanic Garden Jean Massart that was later named after him.

#### Description

Male

Length: body: 1.7 mm; wing: 1.7 mm.

*Head* black in ground-colour. Frons grey dusted, parallel-sided, as wide as pedicel. Face silvery grey dusted, parallel-sided, narrower than pedicel, clypeus shiny black. A pair of long yellowish vertical bristles, widely separated. Antenna entirely black. Postpedicel nearly 2.5 times as long as deep. Arista 2 times as long as all three antennal segments together. Palpus pale yellowish, small, rounded, with a long white apical bristle longer than palpus.

*Thorax*. All bristles yellow. A long humeral, two long notopleurals; acrostichal bristles short, biserial, the rows widely separated. Four long dorsocentrals, the anterior longer than pedicel and bristles becoming longer towards scutellum ending in a long prescutellar; the row is preceded by a few minute bristles. Sternopleura shining black.

*Legs* yellow except for brown mid and hind coxae; all tarsi annulated brown, apical two tarsomeres of all tarsi entirely brown.

Fore coxa with long pale bristles. Fore femur thickened on basal half, wider than mid femur, with a row of long yellow ventral bristles, half as long as femur is wide. Fore tibia hardly spindle-shaped thickened.

Mid femur with a pale brownish anterior bristle on apical quarter, as long as femur is wide. A few yellowish posteroventral bristles, half as long as femur is wide. Mid tibia with a very short, pale brownish apical spur (half as long as tibia is deep), in dorsal view triangular.

Hind femur slender, narrower than mid femur, dorsoventrally bowed, with indistinct bristles.

*Wing* clear, with pale brownish veins, tip of subcostal quite swollen. Veins R4+5 and M parallel, just before ending in the costa, a little diverging. Haltere yellowish white.

*Abdomen* entirely black with short pale bristles. Male terminalia as in Fig. 2. Cerci small enclosed in epandrial lamellae. Right cercus in dorsal view with a broad truncate apex (Fig. 2 B). Right margin of right epandrial lamella with a few of long bristles (Fig. 2 A), near middle of the right lamella a row of 4 short bristles. Left epandrial lamella with marginal bristles on left side very short and limited to apical half (Fig. 2 C); marginal bristles on right side longer than on left side. Inside of apex of left epandrial lamella somewhat sculpted (Fig. 2 B).

#### Female

Length: body: 1.6 mm; wing: 1.4 mm.

Identical to male in most characters including shape and colour of spur on mid tibia. Apical annulation of basal tarsomeres shorter and less pronounced than in male.

#### Comments

*Platypalpus massarti* sp. nov. is found in the samples together with *P. cothurnatus* Macquart, 1827 which is also a small species with entirely black antenna and a small apical spur on the mid tibia. It can be recognized quickly by the coloration of the tarsomeres and the wing. In *P. cothurnatus* only tarsomere 4 and 5 are darkened while the basal tarsomeres are yellow. In *P. massarti* sp. nov. all the tarsomeres of all legs are brown annulated and the apical two tarsomeres entirely brown. In *P. cothurnatus* the wing is conspicuously yellow or yellowish brown clouded while in *P. massarti* sp. nov. there is no yellowish clouding at all. The apical spur in *P. cothurnatus* is nearly as long as tibia is wide and bears a tiny hair at tip in the male. The spur in *P. massarti* sp. nov. is even shorter and lacking a hair. In *P. cothurnatus*, the left epandrial lamella bears a large basal extension on the left side, bearing very long bristles (see Chvála, 1975, fig. 418). In the new species, the marginal bristles on the left epandrial lamella are very short and confined only to the apical half and there is no basal protrusion at all (Fig. 2 C).

A detailed diagnosis of *P. cothurnatus* can be found in Chvála (1975: 166, figs 416-418; 1989: 311).

*P. massarti* sp. nov. belongs to a species-complex comprising *P. cryptospina* (Frey, 1909) and *P. aliterolamellatus* Kovalev, 1971 and hence these two species are dealt with in more detail here below.

### *Platypalpus cryptospina* (Frey, 1909)

Fig. 3

**Fig. 3.**
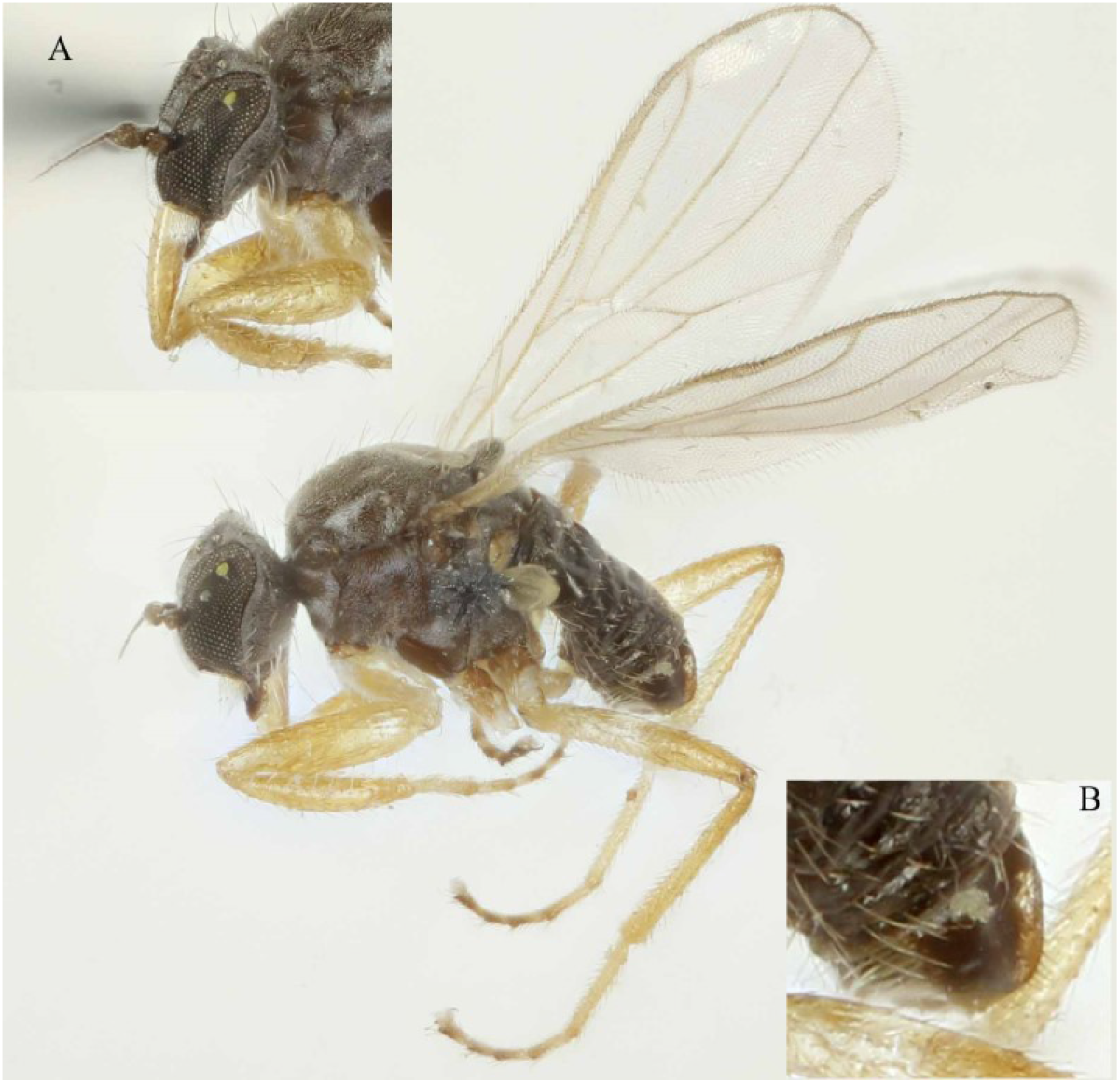
*Platypalpus cryptospina* (Frey, 1909), holotype male habitus; A. detail of head with swollen fore tibiae; B. Left epandrial lamella (photo Jere Kahanpää).

*Tachydromia cryptospina* Frey, 1909: 8.

*Tachydromia tantula* Collin, 1926: 158.

*Tachydromia tantula* in Collin, 1961: 159, re-description and illustration male terminalia (Fig. 59).

*Platypalpus cryptospina* (Frey, 1909) in Chvála, 1975: re-description and illustrations (Figs 125, 216, 419-421, 717).

#### Material examined

Holotype male, Finland, Karislojo, H. Frey, Natural History Museum, Helsinki.

Male on pin, in good condition (missing a mid-leg and left antenna). The male terminalia of the holotype were not dissected so it is unlikely that the drawings of the terminalia in Chvála (1975) were made of the holotype. The drawing of the left epandrial lamella (Chvála, 1975: Fig. 421) fits to the holotype though the long bristles on the left basal 2/3 are even a little longer and the tips are curled and not straight as shown by Chvála (see Fig. 421). In *P. massarti* sp. nov. the third antennal segment is 2.5 times as long as wide and thus longer than in *P. cryptospina. Platypalpus tantulus* (Collin, 1926) was set synonym by Chvála (1975) to *P. cryptospina*, however the bristling on the left side of the left epandrial lamella is different from Chvála’S (1975) drawings and those made by Collin (1961).

To clarify this, the type material of *P. tantulus* should be re-examined. The drawings of the cerci, right and left epandrial lamellae in Collin (1961.) also do not correspond to *P. massarti* sp. nov.

#### Diagnosis

Resembling *P. cothurnatus* but antennae with smaller third segment, dorsocentrals longer and less numerous, legs with dark annulated tarsi, very spindle-shaped fore tibiae and much shorter tibial spur.

### *Platypalpus aliterolamellatus* Kovalev, 1971

*Platypalpus aliterolamellatus* Kovalev, 1971. Description (in Russian); illustration left side of left epandrial lamella.

*Platypalpus aliterolamellatus* in Chvála, 1989: 308, extended diagnosis.

This small species (1.1 – 1.4 mm) is very similar to *P. cryptospina*. The postpedicel is about 1.5 times as long as deep and the arista more than 2 times as long but the acrostichal bristles are wider apart and the legs are paler yellow including the tarsi. At most the two apical tarsomeres are brownish, no annulations are present; the posterior four coxae are yellowish. As in *P. cryptospina* the anterior four femora are equally stout and the fore tibia spindle-shaped dilated. The left border of the left epandrial lamella has very short marginal bristles in the apical third, with some longer bristles in the middle (KOVALEV, 1971: Fig. 1).

#### Key to the *P. cryptospina* complex

1. - Tarsi not annulated brown, at most the apical two tarsomeres brownish; mid and hind coxae not darkened……………*P. aliterolamellatus* Kovalev

-. All tarsi annulated brown, apical two tarsomeres almost entirely brown; mid and hind coxae brownish……………2.

2. - Fore tibia hardly dilated (Fig. 1); third antennal segment in male nearly 2.5 times as long as deep; Acrostichals widely separated. Left epandrial lamella with marginal bristles on left side very short and limited to apical half (Fig. 2 C)……………*P. massarti* sp. nov.

-. Fore tibia distinctly spindle-shaped dilated (Fig. 3); third antennal segment about 2 times as long as deep. Acrostichals close together. Left epandrial lamella with marginal bristles on left side very long over the entire border, tips basal bristles curled (Fig. 3 B)……*P. cryptospina* Frey

### *Platypalpus nanus* (Oldenberg, 1924)

#### Material examined

Oudergem, Botanic Garden Jean Massart: **MT2**, 1 ♀, 9-16.VI.2016; **MT6**, 2♂♂, 3♀♀, 24.V-1.VI.2017; 1♂, 4♀♀, 1-8.VI.2017; 1♀, 8-15.VI.2017.

*Platypalpus nanus* was most abundant in the swampy area (MT6) while only a single female was found during the three-year survey in site 2 (MT2).

This tiny *Platypalpus* species is mainly distributed in coastal areas in Belgium where it was found in nine localities [Blankenberge, Boerekreek, Lombardsijde (Brandaris), Doel, Knokke, Koksijde, Oostende, Raversijde and the Zwin].

It was very abundant in the Maria-Hendrika park in Oostende where a Malaise trap, placed in a wood on clayish soils, collected over two years no less than 363 specimens. Comparably, two Malaise traps placed on wet clayish soil at the base of the dunes of the domain ‘Prince Karel’ at Raversijde, collected over two years 390 specimens. At the moment there are only three observations of *P. nanus* outside the coastal area. A single female was found on a sand heap in the harbour of Ghent (10.VI.1951, leg. M. Bequaert, coll. RBINS), a single female in Brussels (6.VI.1933, leg. M. Bequaert, RBINS) with unknown habitat and another single female was found in an apple orchard in Gembloux (14.VI.1982, leg. C. Fassotte, RBINS). It is not excluded that the latter two females belong to other related unrecognized species.

Chvála (1989) already remarked that the species is very common in Belgium and the Netherlands while he reported it also from the Czech Republic (one record), Germany and Hungary where it is rare. It is not clear which ecological factors explain the distribution of the species in central Europe, while the large abundance in certain coastal areas in Belgium seems to point to a relation with wet, clayish soils.

### *Platypalpus negrobovi* Grootaert, Kustov & Shamshev, 2012 versus *Platypalpus longimanus* (Corti, 1907)

#### *Platypalpus negrobovi* Grootaert, Kustov & Shamshev, 2012

Fig. 4.

**Fig. 4.**
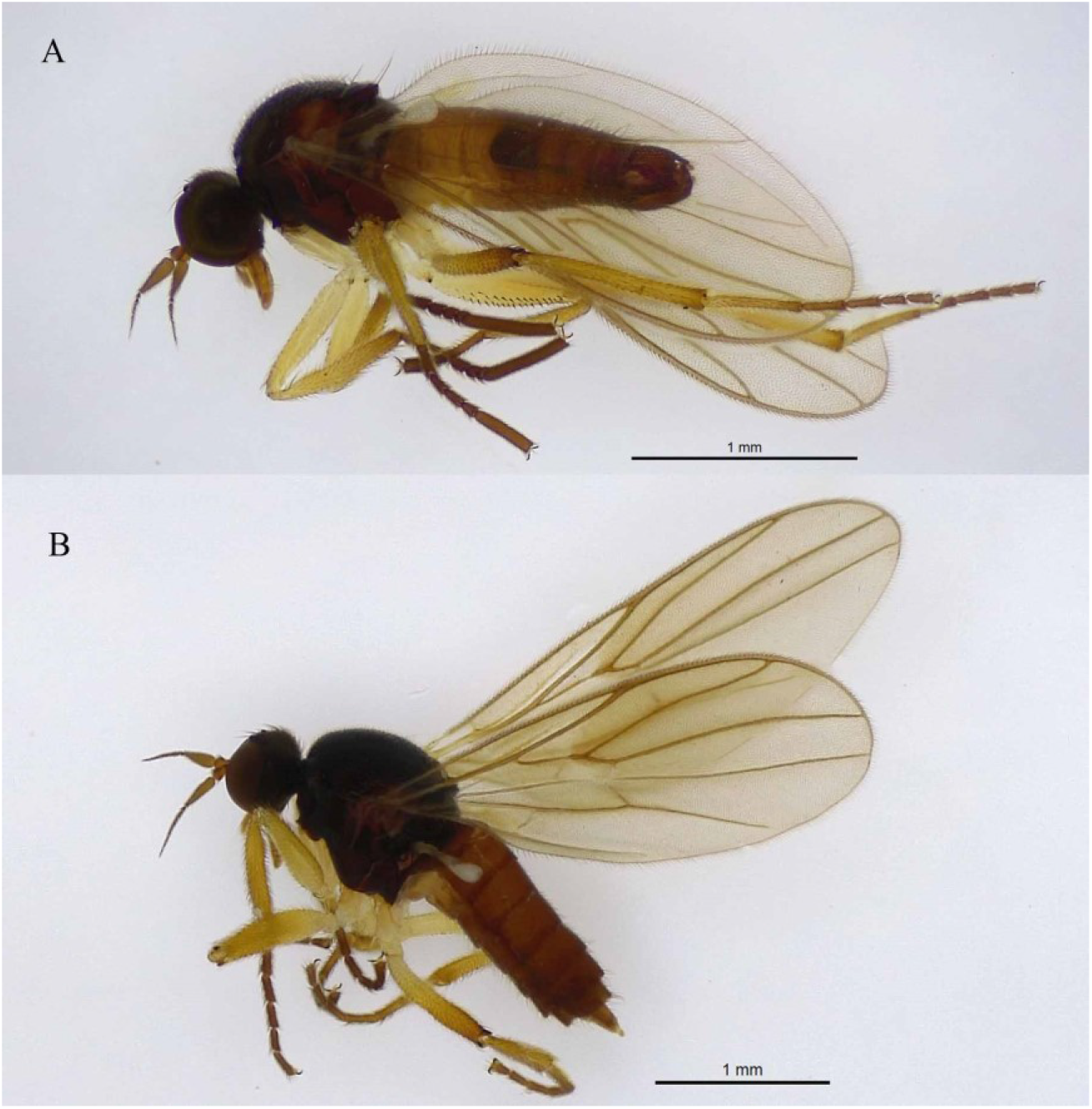
*Platypalpus negrobovi* Grootaert, Kustov & Shamshev, 2012. A, male habitus, Botanic Garden Jean Massart (ref. 1274, RBINS). B, female habitus (photo Isabella Van de Velde).

*Platypalpus negrobovi* Grootaert, Kustov & Shamshev, 2012: 161, illustration antenna (Fig. 1), habitus (Fig. 2), fore and mid leg (Fig. 3), male terminalia (Figs 4–5).

**Fig. 5.**
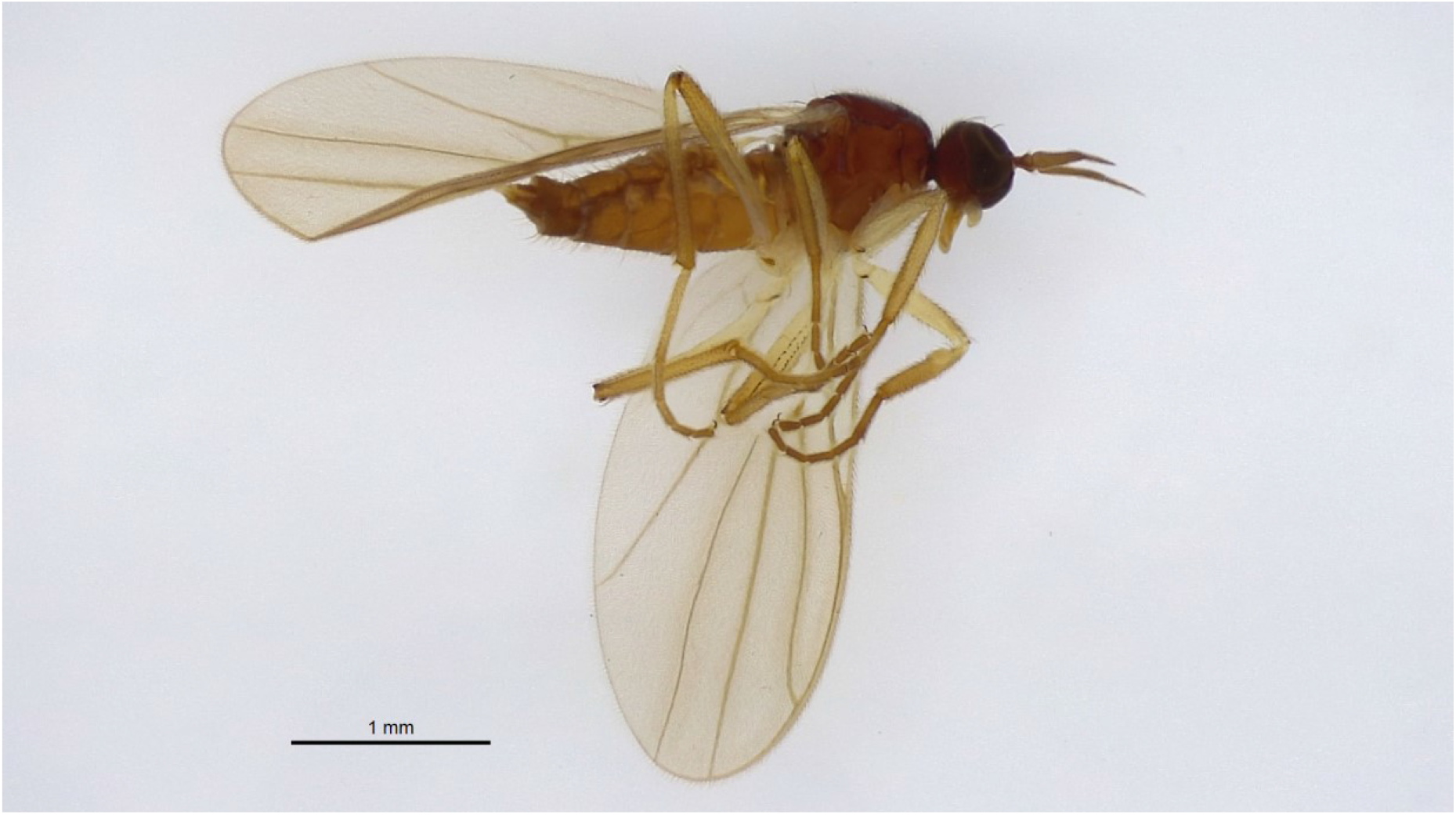
*Platypalpus longimanus* (Corti, 1907), female habitus, Botanic Garden Jean Massart, ref. 1045 (RBINS) (photo Isabella Van de Velde).

##### Material examined

Oudergem, Botanic Garden Jean Massart: **MT2**, 1♀, 9-16.VI.2016 (ref. 892); 1♀, 16-23.VI.2017 (ref. 494); 1♂, 2♀♀, 1-8.VI.2017 (ref. 1274); 2♀♀, 8-15.VI.2017 (ref. 1237); 1♀, 8-15.VI.2017 (ref. 1375); 1♂, 15-22.VI.2017 (ref. 1060); MT5, 2♀♀, 1-8.VI.2017 (ref. 1284); 2♀♀, 22-30.VI.2017; **MT3**, 1♀, 23.VI-1.VII.2016 (ref. 892); **MT5**, 2♀♀, 1-8.VI.2017 (ref. 1284).

*P. negrobovi* was most abundant in site 2 (MT2), but was also recorded in site 3 (MT3) and 5 (MT5).

Males of the sister-species *P. negrobovi* and *P longimanus* are quite peculiar in having a very long flattened apical tarsomere on the fore leg and the mid leg. These are unique features among representatives of the genus *Platypalpus* Macquart, 1827. Smith (1969) gave a re-description of *P. longimanus* and illustrated the fore and mid legs of male as well as the genitalia. There is some confusion about the authorship of *P. longimanus* that has been settled by Chvála (1989). Recently Grootaert *et al*. (2012) described *P. negrobovi* on the base of a single male from the Caucasus although a female was found before in Belgium that was not described by the lack of a male. This female did not correspond to the description of a female *longimanus*.

The differences between the two species are ample discussed in Grootaert *et al*. (2012) and keyed here below.

- Stylus in male nearly as long or a little longer than postpedicel (0.9 – 1.1 times); stylus in female longer: 1.3 to 1.4 times as long as postpedicel. Postpedicel in both sexes shorter than in *P. longimanus*. Male mid tarsus with apical tarsomere as long as tarsomeres 2, 3 and 4 combined.……………*P. negrobovi* Grootaert, Kustov & Shamshev, 2012

- Stylus in male half as long as postpedicel, in female 0.6 – 0.7 times as long. Postpedicel in both sexes longer than in *P. negrobovi*. Mid tarsus with apical tarsomere as long as tarsomeres 3 and 4 together……………*P. longimanus* (Corti, 1907)

In the specimens studied in the present survey, the stylus in males is 1.1 times as long as postpedicel and a little longer in females ranging from 1.3 to 1.4 times as long as postpedicel.

### *Platypalpus longimanus* (Corti, 1907)

Fig. 5

*Tachydromia longimana* Corti, 1907

*Tachydromia longimana* Strobl, 1910

*Platypalpus (Cleptodromia) longimana* (Corti, 1907) in Smith, 1969: 108, illustration fore tibia and tarsus (Fig. 1), tarsus mid leg (Fig. 2), male terminalia (Fig. 3).

*Platypalpus longimanus* in Chvála, 1989: 257, re-description and drawing antenna male (Fig. 3), antenna female (Fig. 4), male terminalia of holotype (Figs 5–7).

#### Material examined

Oudergem, Botanic Garden Jean Massart: **MT2**, 1♀, 11-17.V.2017 (ref. 1046); **MT5**, 1♀, 17-24.V.2017 (ref. 1395).

The postpedicel in female *P. longimanus* is much longer than in *P. negrobovi* and the stylus is much shorter (proportion of 0.65 to 0.7 times as long as stylus).

Identification of both species is sometimes confusing since there are two pairs of black vertical bristles pressed to the cranium, a little longer than the other postocular bristles and pointing forward (not upwards like in other *Platypalpus* with 2 pairs of vertical bristles).

The fore tibiae are swollen in both sexes and the opening of the tibial gland is quite prominent.

### *Platypalpus pictitarsoides* sp. nov

Figs 6–8.

**Fig. 6.**
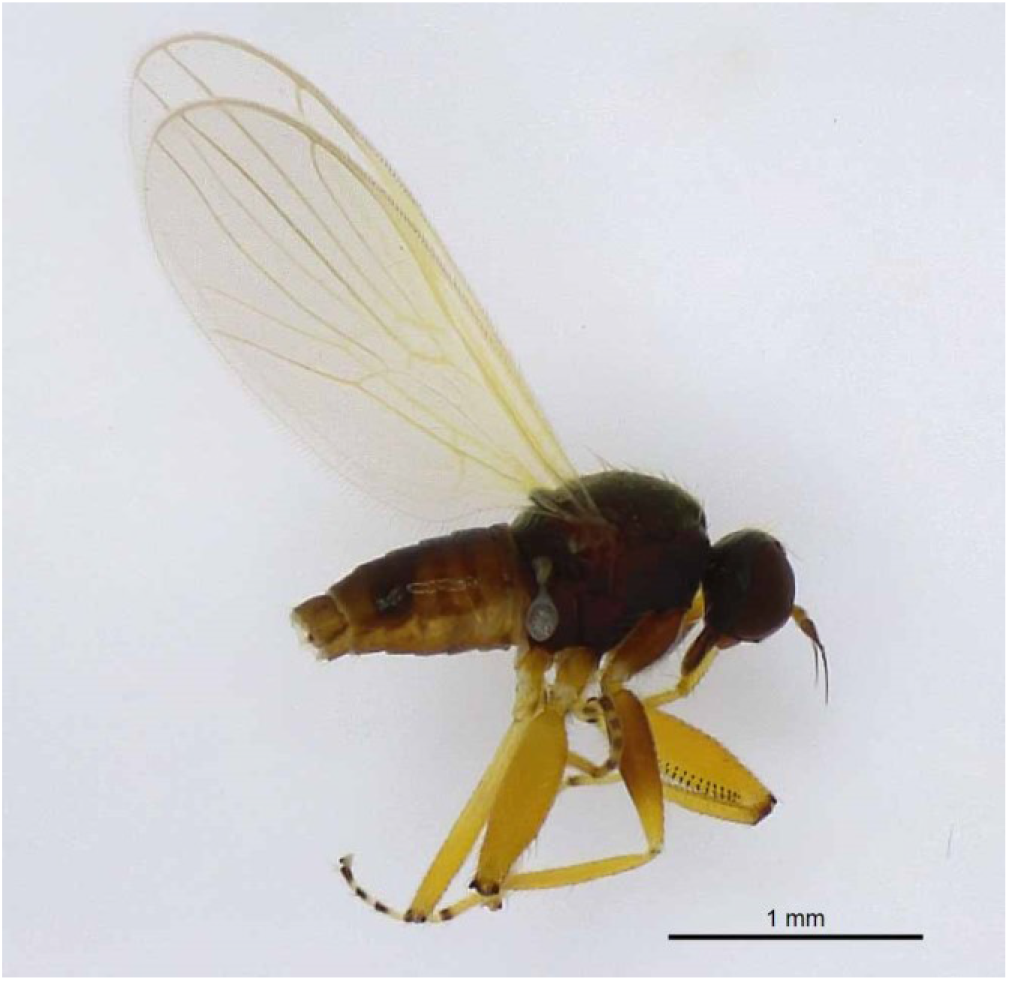
*Platypalpus pictitarsoides* sp. nov., holotype male, habitus (photo Isabella Van de Velde).

**Fig. 7.**
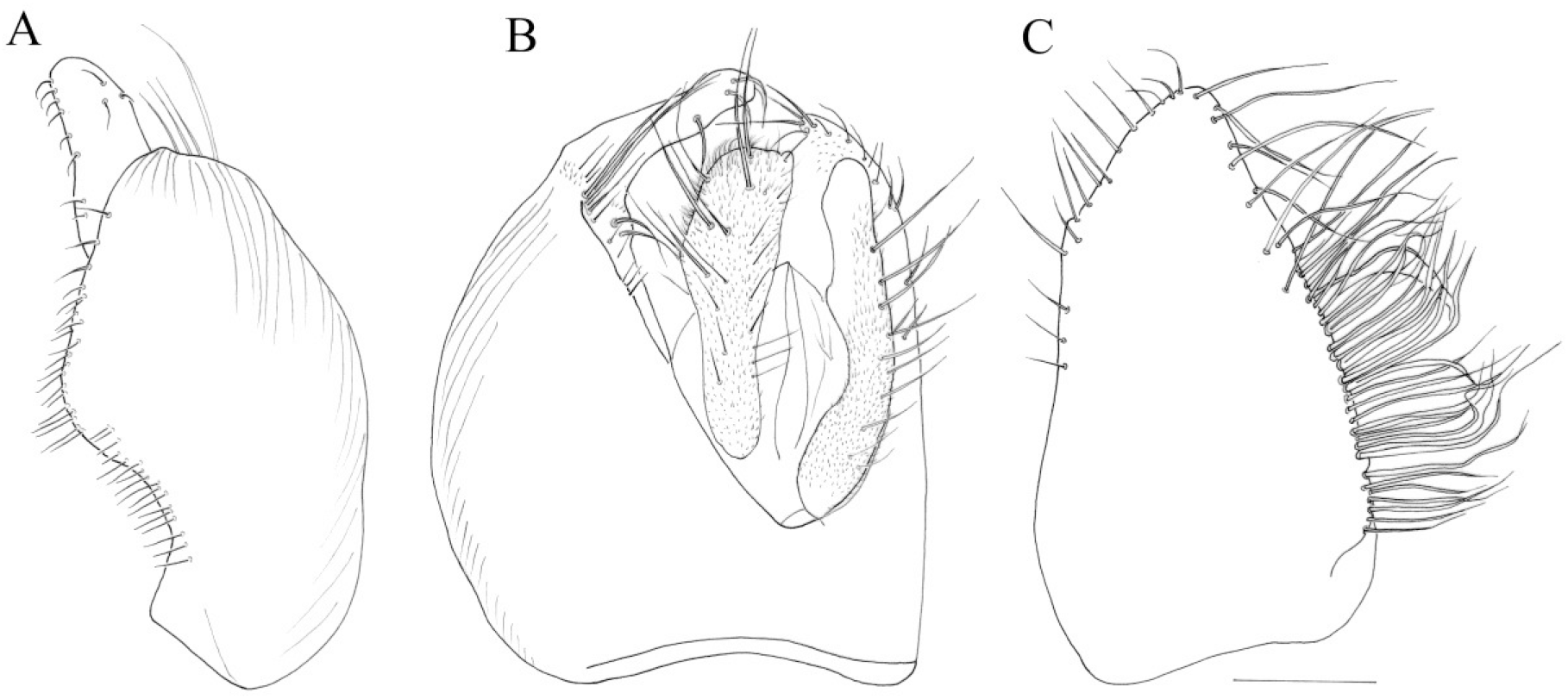
*Platypalpus pictitarsoides* sp. nov., holotype male, terminalia. A, right epandrial lamella; B. epandrium dorsal. C, left epandrial lamella. Scale 0.1 mm.

**Fig. 8.**
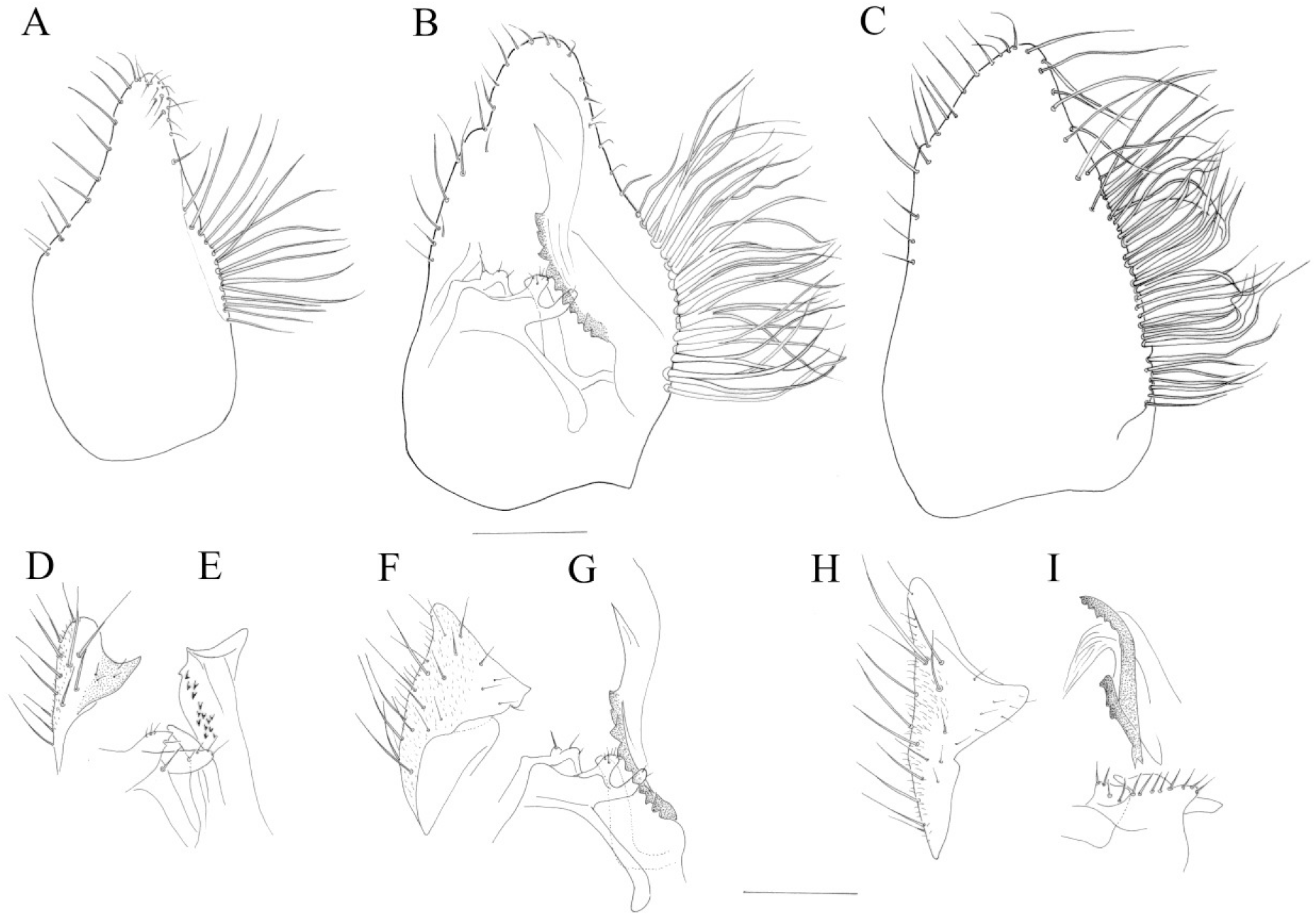
Comparison of left epandrial lamella (A-C), left cercus in lateral view (D, F, H) and aedeagal complex (E, G, I) between *P. kirtlingensis*, *P. pictitarsis* and *P. pictitarsoides* sp. nov. A, *P. kirtlingensis* (paratype, France, St. Cyr en Archies, 1985). B, *P. pictitarsis* (France, St. Cyr en Archies, 1985) with the indication of the position of the aedeagal complex and the inner structures on the left epandrial. C, *P. pictitarsoides* sp. nov. (paratype, oudergem, ref. 1176). D-E, *P. kirtlingensis*. F-G, *P. pictitarsis*. H-I, *P. pictitarsoides* sp. nov. Scale:0.1 mm.

urn:lsid:zoobank.org:act:EA00C554-47D6-4A3F-97EE-F4232A63F336

#### Material examined

Holotype ♂: Belgium, Oudergem, (Brussels) Botanic Garden Jean Massart, **MT6**, 1 – 8.VI.2017 (ref. 1194, RBINS).

Paratypes: Belgium, Oudergem, (Brussels) Botanic Garden Jean Massart, **MT6**, 1♂, 24.V-1. VI.2017 (ref. 1176); 2♀♀ 24.V-1.VI.2017 (ref. 1178); 1♀, 1-8.VI.2017 (ref. 1357); 2♂♂, 9♀♀, 1-8.VI.2017 (ref. 1194); 3♂♂, 42♀♀, 8-15.VI.2017 (ref. 1207); 2♂♂, 46♀♀, 5-22.VI.2017 (ref. 1216); 7♀♀, 22-30.VI. 2017 (ref. 1321); 1♀, 26.VII-2.VIII.2017; 1♂, 1♀, 2-11. VIII.2017; 2♀, 23.VIII-1.IX.2017.

#### Type locality

Audergem; Botanic Garden Jean Massart site 6 (MT6). Only the specimens of site 6, the marshland, were assigned paratype status. In total 201 specimens were found in the Garden, distributed over four sites: MT1: 41, MT2: 30, MT3: none, MT4: none, MT5: 7, MT6: 123.

#### Diagnosis

A small species (1.8 – 2.2 mm) of the *pallidiventris* - *cursitans* group, closely related to *P. pictitarsis*. Clypeus dusted. Third antennal segment 2.5 times as long as deep. Palpus brown in male and female. A very short anterior notopleural present, hardly a third of the length of the 2 posterior notopleurals. Acrostichal bristles biserial, the rows distinctly separated and the bristles directed backward, not uniserial nor diverging. Legs yellow, but fore coxa and basal half of fore femur in male brownish, yellow in female. Mid and hind coxae yellow, at most dusky in male. All tarsomeres annulated black. Mid tibiae in male and female with a long pointed apical spur. Left epandrial lamella with long marginal bristles over the entire length of the outer (left) margin.

#### Etymology

The species is named after its resemblance with *P. pictitarsis*.

#### Description

Male.

Length: body: 1.74 – 1.9 mm: wing: 1.8 mm – 2 mm.

Head. Frons grey dusted, a little narrower than 2^nd^ antennal segment. Face narrower than frons, silvery grey dusted, including clypeus. Basal antennal segments yellowish though somewhat reddish. Third antennal segment black, 2.5 times as long as deep, arista about 1.5 times as long as third segment. A pair of long yellowish white vertical bristles. Lower postoculars bristles longer and whitish.

Thorax grey dusted except for the black shiny spot on sternopleura. All bristles and hairs yellowish. Acrostichals biserial, the rows distinctly separated, the bristles are directed backward and hence not diverging.

Wing with clear membrane, veins pale yellowish. Cross veins well separated. Veins R4+5 and M1 running parallel up to ending in the costa. Haltere white. Squama white, set with white bristles.

Legs yellow except for the brownish fore coxa and basal half of the fore femur that are brownish (the intensity of the browning is variable). Knee fore femur with a very small black spot at both sides of the knee. Mid femur with a large black spot on both sides of the knee. Hind femur with a very small black spot at both sides of the knee. All trochanters with a small black ventral spot. All tarsomeres annulated dark brown.

Fore femur with a double row of white ventral bristles a little more than half as long as femur is wide. Fore tibia tubular, not swollen, with a number of pale brownish dorsal bristly hairs.

Mid femur stronger than fore femur, about 1.5 times wider; with white posteroventral bristles less than half as long as femur is wide. A pale anterior bristle on apical quarter. Mid tibia with a long pointed apical, as long as tibia is deep. Hind femur very narrow with a single row of pale ventral bristles over the entire length, as long as femur is deep. Hind tibia with some dorsal bristly hairs.

Abdomen. All segments brownish black. All tergites with the sides at base narrowly dusted. Male terminalia (Figs 7 – 8). Both cerci are about equally long (Fig. 7 B) The right cercus has a truncate tip while the left cercus is slender with a pointed tip lacking microtrichia at the extreme tip. In lateral view the left cercus bears a large, inner protrusion (Fig. 8 H). The right epandrial lamella has a median protrusion of the right border (Fig. 7 A) and the right border is set with short bristles only. Some striae run over the median part of the right epandrial lamella. The left epandrial lamella is broad and the left margin is entirely set with long bristles. The apical bristles are interspaced while the basal bristles are densely set, some are bifurcate close their base and the tips are often somewhat curled. The aedeagal complex bears two black rod-like and denticulate structures (Fig. 8 I).

#### Female

Length: body: 2 – 2.4 mm, wing: 1.9 – 2.2 mm.

Resembling male in most characters. Legs entirely yellow, including mid and hind coxae. Ovipositor about as long as third antennal segment.

#### Comments

The general morphology of *Platypalpus pictitarsoides* sp. nov. and especially the structure of the male terminalia show that the species is closely related to *P. pictitarsis* and *P. kirtlingensis* (Fig. 8 A, D-E). However, *P. pictitarsoides* sp. nov. has the rows of biserial acrostichals well separated and not on almost one line with diverging bristles like in the former species. The new species has only a tiny anterior notopleural bristle in front of the longer posterior notopleurals bristles which can easily be overlooked or considered as simple pubescence. This may lead to some confusion when using the key of Grootaert & Chvála (1992).

The presence of a tiny anterior notopleural bristle and the fore and hind tibiae with distinct bristly dorsal hairs brings the new species to couplet 141 in the key Grootaert & Chvála (1992). The following key is simplified.

1.- Anterior notopleural bristle strong, nearly as long as posterior notopleurals. Ovipositor much longer than all antennal segments together……*P. pallidiventris* (Meigen) and *P. longiseta* (Zetterstedt)

-. Anterior notopleural bristle smaller and finer, at most half as long as posterior notopleural bristles. Ovipositor about as long as all antennal segments together……………2.

2. - The rows of acrostichal bristles distinctly separated and directed backward and hence not diverging. Left epandrial lamella with a row of long bristles on the entire outer (left) margin (Fig. 7 C; Fig. 8 C)……………*P. pictitarsoides* sp. nov.

-. Acrostichals bristles closely bi-serial, almost alternating on one row and strongly diverging. Left epandrial lamella with short marginal bristles on apical part while the long bristles are confined to the central part of the outer margin (Fig. 8 A, B)……………3.

3. - All coxae and trochanters black (fore coxa apically pale in female); fore femur more or less blackish on basal half, tarsi with sharp black annulations. Palpus brownish-black in both sexes. Left epandrial lamella rather blunt at tip and wider (Fig. 8 B), outer (left) side with very long, bifurcated bristles……………*P. pictitarsis* (Becker)

-. Legs yellow including all coxae and the annulated black tarsi. Palpus yellow in male, dusky in female. Left epandrial lamella apically narrowed and rather pointed, outer (left) margin with less numerous, shorter and simple bristles (Fig. 8 A)……………*P. kirtlingensis* Grootaert

## Comparison of the male terminalia in *P. kirtlingensis*, *P. pictitarsis*, *P. pictitarsoides* sp. nov

(Fig. 8).

Without preparation of the male terminalia, the three species can already be recognized by the shape of the left epandrial lamella. The apex of the left epandrial lamella is quite slender in *P. kirtlingensis* while broader in *P. pictitarsis* and even broader in *P. pictitarsoides*. In *P. pictitarsis* the long marginal bristles on the left side are confined to the basal half, while in *P. pictitarsoides* even on the apical half there are very long marginal bristles present, but more interspaced than at the base. In *P. kirtlingensis* there is a bundle of long bristles, less dense than in the other two species. Moreover, the bristles are simple and not forked.

The tip of the left cercus, more particularly the inward directed apical protrusion is also different. Removal and clearing of the terminalia is generally needed to observe this structure that is laying below the left epandrial lamella in lateral view. In *P. kirtlingensis* (Fig. 8 D) the protrusion is pointed, brownish and directed upward. In *P. pictitarsis* the tip of the left cercus resembles a cap (Fig. 8 G). In *P. pictitarsoides*, the tip of the left cercus is elongated (Fig. 7 B, dorsal view) and lacks microtrichia at the extreme tip (Fig. 8 H). The inward directed protrusion has a different shape than the two other species (Fig. 8 H).

The aedeagal complex is also different. The position of the different parts is illustrated in a lateral view Fig. 8 B, beneath the left epandrial lamella. In *P. kirtlingensis* (Fig. 8 D) the denticulated part consist of a few scattered downward directed denticles. The denticles are more robust and inserted on a dark sclerotized sclerite in *P. pictitarsis*, while in *P. pictitarsoides* there are two separated denticulate sclerotizations, with denticles less strong than in *P. pictitarsis*.

The inner structures on the left epandrial lamella are not well understood. However, there are many differences between the different species. In *P. pictitarsis* the transverse rod-like structure ends in fork each bearing a few tiny apical hairs (Fig. 8 G). In *P. pictitarsoides* this rod-like structure is apically not forked and bears a row of hairs all along the apical side (Fig. 8 I). The structure of the transverse rod-like structure is comparable in *P. kirtlingensis* though the apical row of hairs is limited to a few scattered hairs (Fig. 8 E).

## Comparison with *P. stabilis* (Collin, 1961) and *P. annulitarsis* Kovalev, 1978

Both *P. stabilis* and *P. annulitarsis* lack an anterior notopleural bristle, but have similar male terminalia as in *P. pictitarsoides*. The male terminalia of *P. stabilis* (Collin, 1961) do resemble somewhat those of *P. pictitarsoides* in that the outer marginal bristles of the left epandrial lamella are long along the entire margin. However, the tarsi in *P. stabilis* are faintly brownish annulated and darker on the apical two tarsomeres. In *P. pictitarsoides* they are equally annulated black. The third antennal segment is about twice as long as deep, black with yellow base. The third antennal segment is 2.5 times as long as deep and entirely black in *P. pictitarsoides*.

*Platypalpus annulitarsis* Kovalev, 1978 is also a small species (less than 2 mm in body length) that resembles *P. pictitarsoides*. It has a polished clypeus. The apical half of the third antennal segment is black and the base yellow. The male has a long, blunt-tipped apical spur on the mid tibia which is pointed in the female. In *P. pictitarsoides* the mid tibial spur is pointed in both sexes. The left epandrial lamella is indented on its outer margin and the basal part is protruding with several rows of very long bristles. The outer margin of the left epandrial lamella is not indented and the basal part is not protruding in *P. pictitarsoides*. In addition, the right epandrial lamella is on the basal half of the right (outer) margin set with long bristles. In *P. pictitarsoides* there are only very short bristles all along the outer margin (Fig. 7 A).

### *Platypalpus pictitarsis* (Becker, 1902)

Fig. 8 B, F, G.

*Platypalpus ruficornis* Macquart, 1850 (non von Roser, 1840): 97.

*Tachydromia pictitarsis* Becker, 1902: 44.

*Tachydromia pictitarsis* Becker, 1902 in COLLIN, 1961: 168, re-description and illustration male terminalia (Fig. 62).

*Platypalpus pictitarsis* (Becker, 1902) in GROOTAERT, 1986: 190, re-description and illustration antenna, fore leg, male terminalia (Figs 8-13).

*Platypalpus pictitarsis* (Becker, 1902) in Chvála, 1989: 349, diagnosis.

*Platypalpus pictitarsis* (Becker, 1902) in GROOTAERT & Chvála, 1992: 175, diagnosis and illustration antenna, fore leg, male terminalia (Figs 190-195).

As can be seen in the synonym list above, *Platypalpus pictitarsis* was several times subject of discussion. COLLIN (1961) clearly indicated variability of the characters which led to the later description of *P. kirtlingensis* Grootaert, 1986. COLLIN (l.c.) was hesitative of the distinction of the two species and in fact he illustrated the left epandrial lamella (Fig. 62) from *P. kirtlingensis* while the description is from *P. pictitarsis*. DNA barcoding might be a useful tool to elucidate the genetic differences hopefully corroborating the morpho-species as we identify them now.

### *Platypalpus rapidoides* Chvála, 1975

#### Material examined

Oudergem, Botanic Garden Jean Massart: **MT1**: 1♂, 1♀, 23.VI-1.VII.2016 (ref. 1316); **MT2**: 1♀, 21-28.V.2015 (ref. 623); 1♀, 4-10.VI.2015 (ref. 340); **MT6**: 1♂, 1-8.VI.2017 (ref. 1197).

A rare species often found together with *P. rapidus* (Meigen). *P. rapidus* has the fore coxa and femur yellow, while they are black in *P. rapidoides*. The acrostichals are quadri-serial in *P. rapidus* while irregularly six-serial in *P. rapidoides*.

### *Platypalpus subtilis* (Collin, 1926)

#### Material examined

Oudergem, Botanic Garden Jean Massart: **MT1**: 1♀, 6-12.VIII.2015 (ref. 131); **MT2**: 1♀, 28.VII-4.VIII.2016 (ref. 980); **MT3**: 1♀, 1-6.VII.2016; 1♀ (ref. 945); 1♀, 6-14.VII.2016; **MT5**: 1♀, 30.VI-6.VII.2017 (ref. 1330); **MT6**: 1♀, 24.V-1.VI.2017 (ref. 1181); 1♀, 8-15.VI.2017 (ref. 1205).

Additonal material examined: Belgium: 1♀, Gomery, 21.V.1981; 1♀, Ottiginies, 1.VIII.1981; 1♀, 15.VIII.1981; 1♀, 29.VIII.1981; 1♀, Buzenol, 16.VI.1981; 1♀, 30.VI.1981; 1♀, 28.VII.1981; 1♀ 11.VIII.1981.

An enigmatic species known from only seven females in the Botanic Garden. The eight earlier records of this species from the South of Belgium are also females. Probably confused with other species. To our knowledge, there are no figures of the male terminalia available.

### *Trichina opaca* Loew, 1864

*Trichina opaca* Loew, 1864: 40.

*Trichina picipes* Tuomikoski, 1935: 99.

*Trichina opaca* Loew, 1864 in Chvála, 1983: 130 and illustrations (Figs 231, 232, 249-254).

#### Material examined

Oudergem, Botanic Garden Jean Massart: **MT2**, 1♀, 8-15.VI.2017 (ref. 1201).

#### Comments

Using the key of Chvála (1983), the female found here at the Botanic Garden leads to *T. opaca* in having the anterior part of the mesoscutum covered with microtrichia. The third antennal segment is broader and much shorter than in *T. elongata* and the stylus is about half as long as the third segment. According to Chvála (1.c.) the female is much paler than the male which would fit to the female found here. Males are needed to confirm the presence of this species in Belgium.

#### Distribution

According to the Fauna Europaea, *T. opaca* is recorded from the British Isles, Central European Russia, Czech Republic, Finland, Germany, Ireland, Italy and Switzerland. Its occurrence in Belgium was thus expected.

### *Stilpon subnubilus* Chvála, 1988

#### Material examined

Oudergem, Botanic Garden Jean Massart: **MT2**, 1♂, 12-20.VIII.2015; 1♀, 24.XI-20.XII.2016 (ref. 401).

The species was recently reported from The Netherlands (Belgers *et al*., 2021). It is very closely related to *S. moroccensis* Grootaert *et al*., 2021.

## Discussion

In the present three-year survey of the Botanic Garden Jean Massart, 90 hybotid species were found, representing about 52 % of the species ever recorded in Belgium (Grootaert, 1991; Grootaert *et al*., 2001).

The species accumulation curve (Fig. 9) calculated with the programme EstimateS (Colwell (2013) shows the relation between number of sampling sites/seasons and number of species. The total number of species observed is 90 while the predictions Chao1 predicts 99 species, Bootstrap 97 species and Jackknife 103 species. The number of singletons and doubletons are clearly descending.

**Fig. 9.**
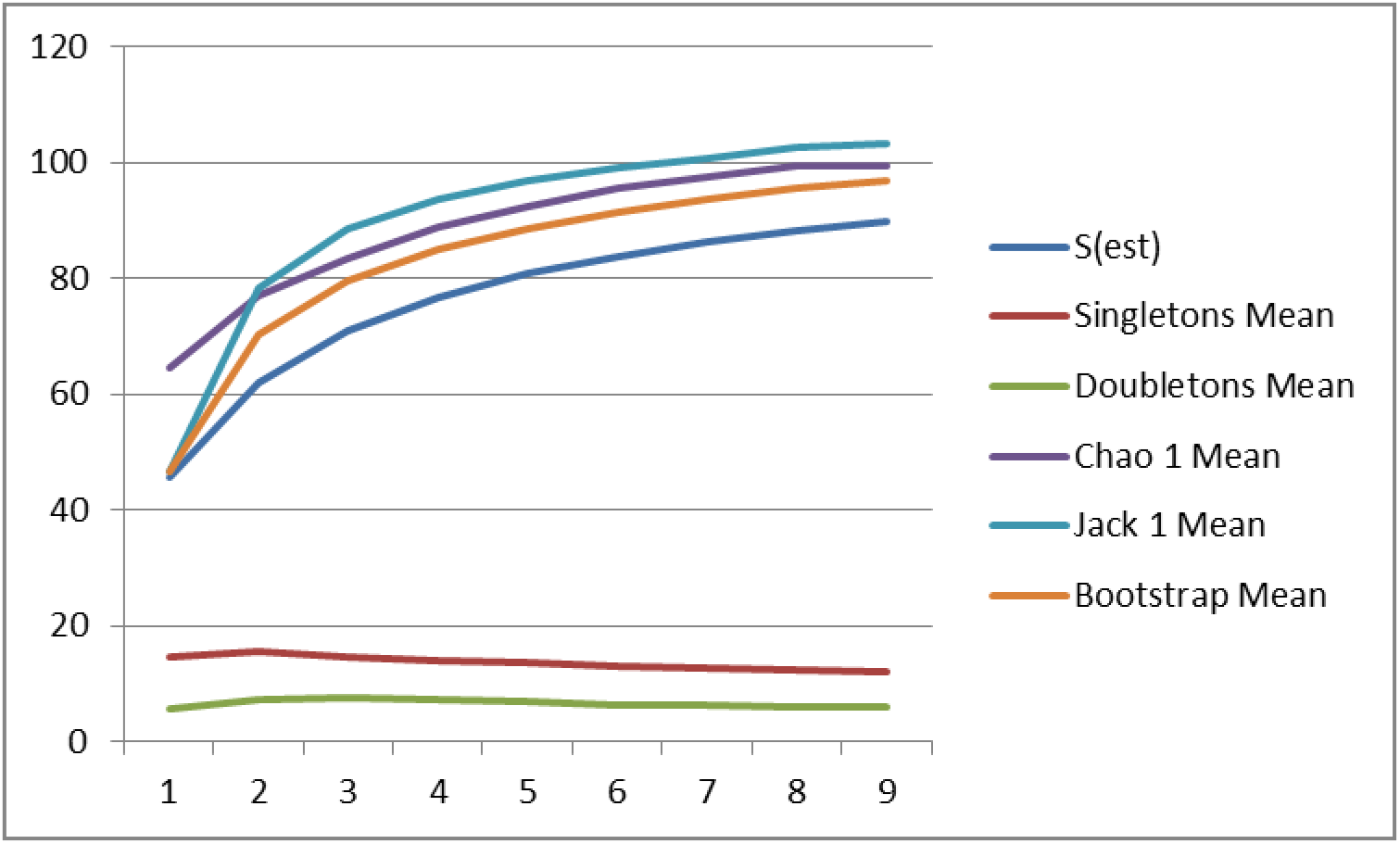
Species accumulation curve calculated with the programme EstimateS (Colwell (2013). Relation between number of sampling sites/seasons (9) and number of species.

### Analysis of the RDB data

Annexe 2 shows the alphabetic list of the hybotid species found at the Botanic Garden Jean Massart with their Red Data Book status, while Annexe 3 shows the list of the Red Data Book status with the species assigned to them from highly threatened species to Safe/Low risks species. Table 2 gives a summary of the number of species per RDB category (Grootaert *et al*., 2001).

**Table 2.** Summary of the number of species per RDB category.

| **RDB category** | **nr species** |
| --- | --- |
| 0. Extinct | 2 |
| 1. Critically endangered | 0 |
| 2. Endangered | 2 |
| 3. Vulnerable | 1 |
| 4. Susceptible | 34 |
| 5. Near threatened | 1 |
| 6. Safe/Low risk | 32 |
| 7. Data deficient | 18 |

Two species that were supposed to be ‘Extinct’ in Flanders and Brussels, were recorded here in the Botanic Garden: *Platypalpus pseudorapidus* Kovalev, 1971 and *Trichina bilobata* Collin, 1926. It is clear that these two species should be removed from this category and be placed in the category 1 ‘Critically endangered’ species.

No species belonging to the category ‘Critically endangered’ were found, while two species belong to the category ‘Endangered’ and one species is considered ‘Vulnerable’. On the other hand, one third of the species, more precisely 34 species, are considered as ‘Susceptible’. This category groups species that are uncommon but for which no significant recent decrease in geographical distribution could be demonstrated. It also groups species from rare habitats.

Another third of the species (32 species) are considered to be ‘Safe/Low risk’. Finally, 18 species are considered as ‘Data deficient’ which means that no trend of decline is visible, simply because they have not been observed before the pivot date of 1980 and hence the percentage observed before and after 1980 cannot be calculated. In addition, these species were not observed in threatened habitats which would put them otherwise automatically in a threatened category. In reality, most of these species are rare and even when more surveys would be available, it is likely that they would not belong to the ‘ Safe/Low risk’ category.

Since its publication in 2001, the Red Data Book on empidids (Grootaert *et al*., 2001) has not been updated nor the data collected since 2001 up to present have been re-calculated nor re-evaluated. Nevertheless, the low percentage of 30% ‘Safe/Low risk’ species is an indication that there is a high number species in the Botanic Garden that are in one way or another threatened species. We are well aware of the main criticisms of Red Data Books that sampling efforts are never sufficient and that they do not reflect the exact condition of the population. According to Heijerman & Turin (1998), the results are therefore arbitrary. These arguments are valid but it must also be recognised that there is an urgent need for properly processed data, and that such data, even if incomplete, may reveal real patterns. Site quality assessments are preferably based on a combination of species’ richness, abundance, rarity and vulnerability estimates of as many biota as possible. As reliable data on rarity and vulnerability are almost entirely restricted to Red Data Books, it seems evident that only organisms which have been investigated in this respect can be used in these assessment studies.

## Acknowledgements

The present paper is an output of the project ‘Objective 1000’, commissioned by Brussels Environment. Sincere thanks go to Mr Alain Drumont and Mr Hugo Raemdonck (RBINS) for the weekly collecting of the samples. The enthusiastic and very helpful support of the staff of the Botanic Garden Jean Massart, especially the help of Mr Thierry Bruffaerts, is also much appreciated. Isabella Van de Velde (RBINS) and Jere Kahanpää (LUOMUS) helped with the photos.

---

**Annexe 1.**
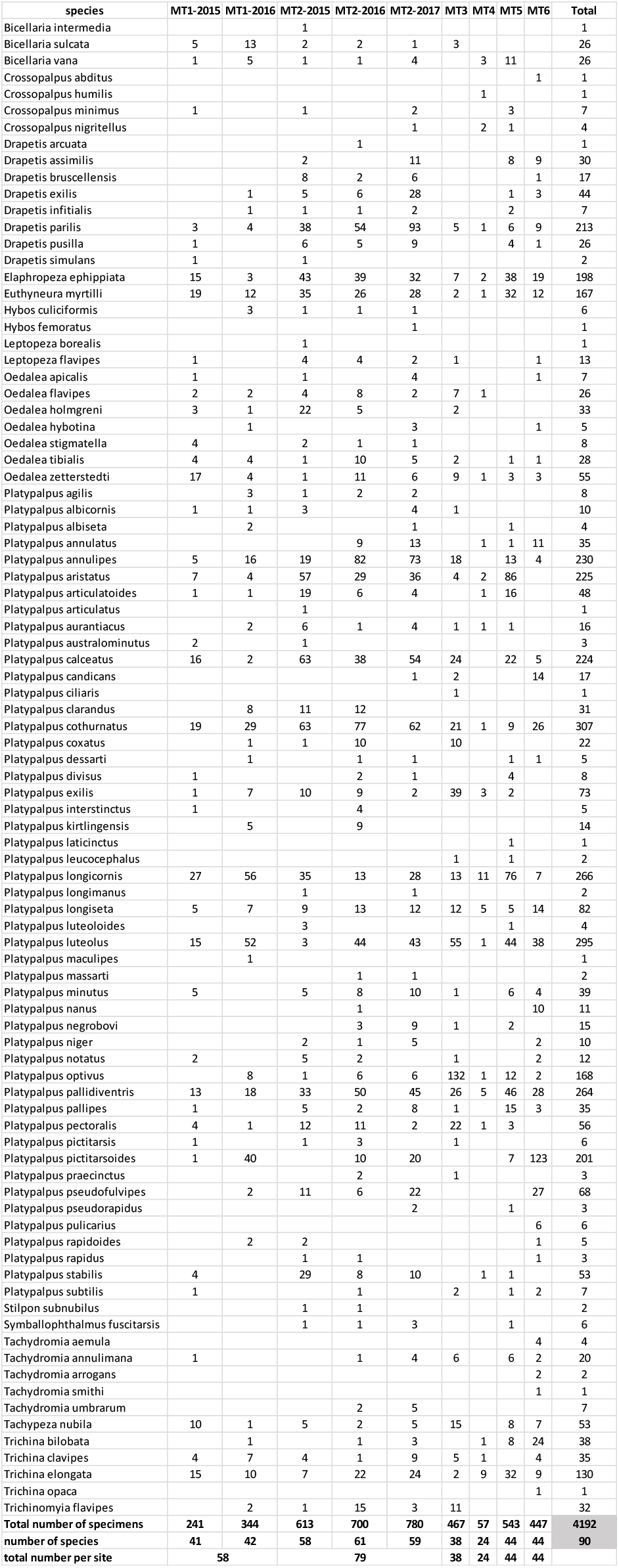
Annexe 1. Overview of the number of species per site/per year. Site 1 (MT1) and site 2 (MT2) were respectively sampled during 2 and 3 years, the data are presented for each year separately (MT1-2015, MT1-2016; MT1-2015, MT1-2016 and MT1-2017). Site 3 (MT3) was only sampled in 2016 while sites 4, 5 and 6 (MT4, MT5 and MT6) were only sampled in 2017.

**Annexe 2.**
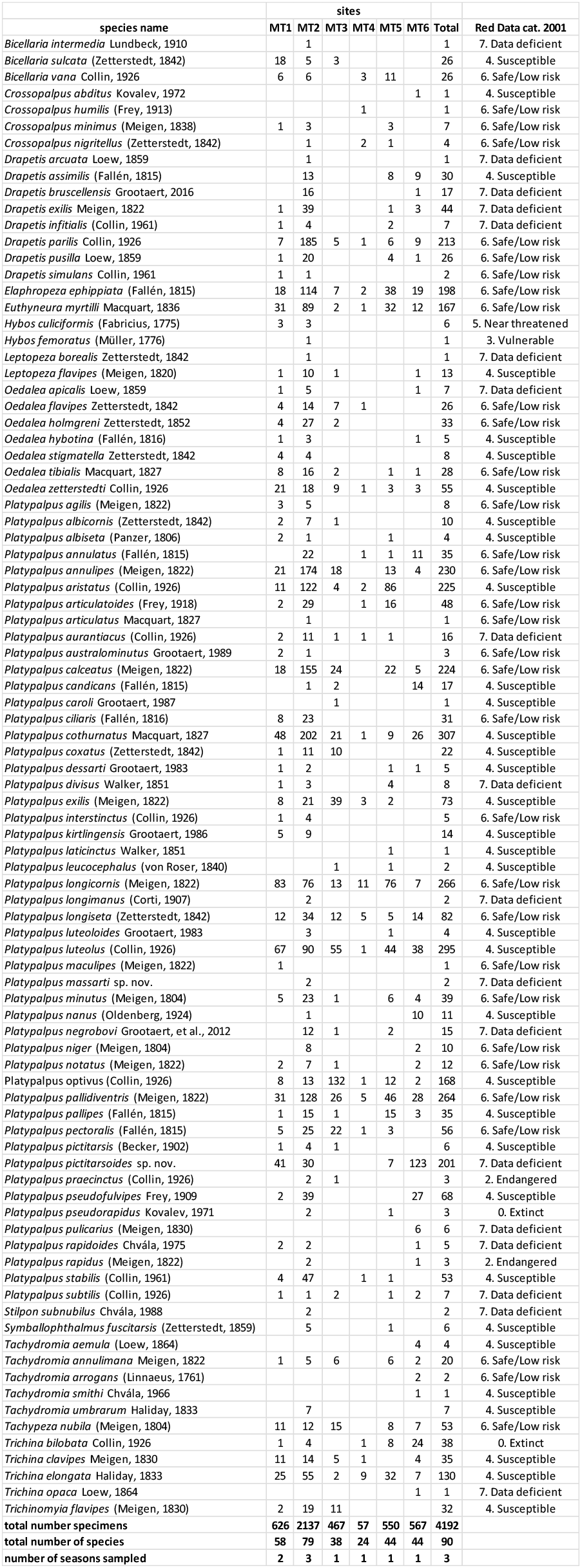
Annexe 2. Alphabetic list of the hybotid species found at the Botanic Garden Jean Massart with their Red Data Book status.

**Annexe 3.**
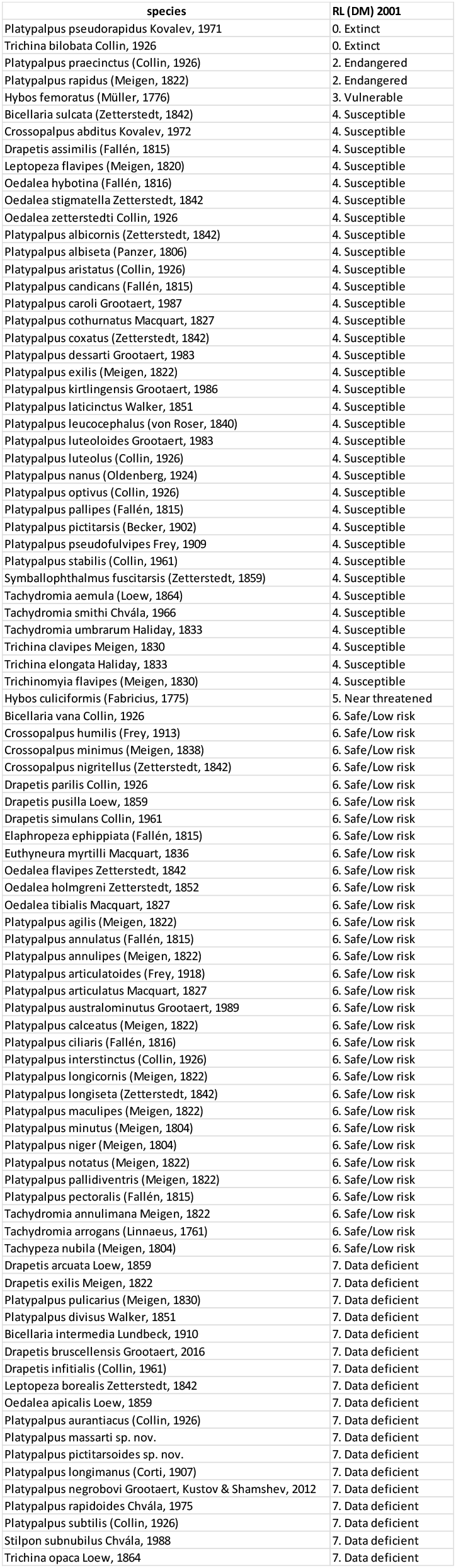
Annexe 3. List of the Red Data Book status with the species assigned to them.

## References

Belgers J.D.M., Beuk P., van der Weele R. & Prijs J., 2021. - *Stilpon subnubilus:* een nieuwe bocheldansvlieg voor Nederland (Diptera: Hybotidae). Entomologische Berichten, 81: 21–24.

Chvála M., 1975. - The Tachydromiinae (Dipt. Empididae) of Fennoscandia and Denmark. Fauna entomologica Scandinavica, 3: 336 pp. Klampenborg, Denmark.

Chvála M., 1983. - The Empidoidea (Diptera) of Fennoscandia and Denmark. II. General Part. The families Hybotidae, Atelestidae and Microphoridae. Fauna entomologica scandinavica, 12: 279 pp.

Chvála M., 1988. - A new species of *Stilpon* loew (Dipt., Hybotidae) related to *S. nubilus* Collin from England and western Europe. Entomologist’s monthly magazine, 124: 255–231.

Chvála M., 1989. - Monograph of northern and central European species of *Platypalpus* (Diptera, Hybotidae), with data on the occurrence in Czechoslovakia. Acta Universitatis Carolinae - Biologica, 32: 209–376.

Collin J.E., 1961. - Empididae. In: British Flies. Vol. 6. University Press, Cambridge. 782 pp.

Colwell R.K., 2013. - EstimateS: Statistical estimation of species richness and shared species from samples. Version 9. http://purl.oclc.org/estimates (Accessed 22 November 2013).

Grootaert P., 2016. - *Drapetis bruscellensis* (Diptera: Hybotidae) a new species for science from the outskirts of Brussels, a not so cryptic species supported by COI barcoding. Belgian Journal of Entomology, 41: 1–14.

Grootaert P. & Chvála M., 1992. - Monograph of the genus *Platypalpus* (Diptera: Empidoidea, Hybotidae) of the Mediterranean region and the Canary Islands. Acta Universitatis Carolinae - Biologica, 36 (1-2): 226 pp, 262 figs.

Grootaert P., Kustov S.Yu. & Shamshev, I.V., 2012. - *Platypalpus negrobovi* a new species of the family Hybotidae (Diptera: Empidoidea) from the North-West Caucasus. Caucasian entomological Bulletin, 8: 161–163.

Grootaert P., Pollet M. & Maes D., 2001. - A Red Data Book of empidid flies of Flanders (northern Belgium) (Diptera, Empididae s.1.): Constraints and possible use in nature conservation. Journal of Insect Conservation, 5: 117–129.

Grootaert P., Zouhair L. & Kettani K., 2021. - A new species of the genus *Stilpon* Loew, 1859 from Morocco (Diptera: Empidoidea, Hybotidae). Belgian Journal of Entomology, 113: 112.

Heijerman T. & Turin H., 1998. - The Red Data Lists: sense or nonsense? Entomologische Berichten, 58: 92–104.

Kovalev V.G., 1971. - New European species of the genus Platypalpus Macq. (Diptera, Empididae) Entologischeskoe Obozrenie, 50: 200–213.

Smith K.V.G., 1969. - *Platypalpus (Cleptodromia) longimana* Corti, New to Britain and the male of *P. altera* (Collin) (Dipt., Empididae). Entomologist’s Monthly Magazine, 105: 108–110.

